# Circadian rhythms mediate malaria transmission potential

**DOI:** 10.1101/2024.05.14.594221

**Authors:** Inês Bento, Brianna Parrington, Rushlenne Pascual, Alexander S Goldberg, Eileen Wang, Hani Liu, Mira Zelle, Joseph S Takahashi, Joshua E Elias, Maria M Mota, Filipa Rijo-Ferreira

## Abstract

Malaria transmission begins when infected female *Anopheles* mosquitos deposit *Plasmodium* parasites into the mammalian host’s skin during a bloodmeal. The salivary gland-resident sporozoite parasites migrate to the bloodstream, subsequently invading and replicating within hepatocytes. As *Anopheles* mosquitos are more active at night, with a 24-hour rhythm, we investigated whether their salivary glands are under circadian control, anticipating bloodmeals and modulating sporozoite biology for host encounters. Here we show that approximately half of the mosquito salivary gland transcriptome, particularly genes essential for efficient bloodmeals such as anti-blood clotting factors, exhibits circadian rhythmic expression. Furthermore, we demonstrate that mosquitoes prefer to feed during nighttime, with the amount of blood ingested varying cyclically throughout the day. Notably, we show a substantial subset of the sporozoite transcriptome cycling throughout the day. These include genes involved in parasite motility, potentially modulating the ability to initiate infection at different times of day. Thus, although sporozoites are typically considered quiescent, our results demonstrate their transcriptional activity, revealing robust daily rhythms of gene expression. Our findings suggest a circadian evolutionary relationship between the vector, parasite and mammalian host that together modulate malaria transmission.

## Introduction

Malaria, a life-threatening disease transmitted by female *Anopheles* mosquitos, continues to pose a significant global health and economic burden ^1^. Malaria transmission is dependent on a mosquito’s behavior of seeking a bloodmeal, which predominantly occurs at night ^2–8^. This circadian-driven behavior allows infected mosquitos to deposit *Plasmodium* parasites into the skin of a mammalian host at a specific time of the day. During a blood meal, uninfected mosquitos can also ingest transmissible parasite forms (gametocytes) from an infected host, which upon ingestion, undergo sexual reproduction forming oocysts in the mosquito’s midgut, eventually giving rise to thousands of motile sporozoites. The sporozoites then travel through the mosquito’s hemolymph, crossing cell membranes until they reach the salivary glands ^9,10^, where they reside for several days ready to encounter another mammalian host and establish a new infection.

While probing the skin for a bloodmeal, female mosquitos inject saliva into the host dermis. Their saliva modulates host responses, including platelet aggregation, coagulation, thrombin activation, vasodilation, and even inflammatory response ^11–13^. Multiple proteins produced in the salivary glands are secreted in the saliva, such as *Anopheles*-specific SG1 (salivary gland 1) gene family (including TRIO ^14^), apyrase ^15^ and anopheline antiplatelet protein (AAPP) ^16^. These saliva proteins have been shown to be important for efficient bloodmeals ^15,17^. Interestingly, saliva proteins can impact parasite transmission, one example is by increasing parasite load during infection ^18,19^. Saliva ingested together with blood also modulates parasite development in the midgut of the mosquito ^15,20^. Additionally, antiserum against salivary gland extracts has also been shown to decrease *Plasmodium* infection in rodent models ^21^. Parasites may modulate vector physiology during infection, as some saliva proteins are expressed at higher levels in infected mosquitoes than in uninfected ones ^13,22^. Together, these studies highlight how understanding salivary gland biology may promote development of novel strategies for controlling this infection.

Various mammalian tissues exhibit daily rhythmic gene expression that regulates physiological functions ^23–26^. Similarly, despite a lack of tissue resolution, *Anopheles* mosquitos were shown to possess transcripts with circadian rhythmic expressions ^27,28^, controlled by core clock genes such as *period* (*per*), *timeless* (*tim*), and *clock* (*clk*). These clock genes, along with environmental cues, coordinate *Anopheles* pheromone synthesis, swarming, and mating behaviors ^29^. Alongside other molecular components, these clock genes are likely to maintain the robustness and precision of the mosquito’s internal clock, ultimately influencing its feeding time behavior and potentially the transmission of malaria parasites. However, it is unknown whether mosquito salivary glands are under circadian control and whether these rhythms influence the daily biology of salivary gland sporozoites. Considering female mosquitos’ cyclic feeding behavior and the importance of the salivary glands during bloodmeals, we hypothesize that a subset of salivary gland genes would be transcribed every 24 hours. This circadian rhythmic expression profile would allow the anticipation and preparation for bloodmeals. Sporozoites are exposed to complex signals within the mosquito’s salivary glands ^30^, where changes throughout the day might influence not only their biological timing, but also potentially impact parasite transmission and, consequently, disease outcome.

In this study, we unveiled the dynamic nature of the mosquito’s salivary gland transcriptome and its relationship with the circadian clock. Our transcriptomic analyses identified that a significant proportion of genes involved in blood-feeding-related functions (including anti-blood clotting) exhibit circadian rhythms of expression in the salivary glands. Moreover, we found that salivary gland sporozoites also have a daily rhythmic transcriptome, particularly concerning genes associated with parasite transmission. These findings suggest cyclic molecular changes in both the mosquito and the parasites that prepare them for bloodmeals. We propose that the interaction of host, parasite and vector circadian clocks is critical for efficient *Plasmodium* transmission. Understanding the molecular mechanisms underlying the intricate interplay between mosquito vector, *Plasmodium* parasite, and mammalian host, will allow us to gain a deeper insight into the biology of malaria transmission, holding an immense potential for the development of novel intervention strategies. These findings may extend to other infectious diseases that cause major public health burdens, such as Zika and dengue that are also vector-borne and thus may have daily rhythms in transmission. Ultimately, this will contribute to the global effort in the combat against these devastating infectious diseases.

## Results

### Mosquito salivary glands have circadian rhythms that correlate with mosquitos’ feeding behavior

Despite evidence that mosquitos have daily rhythms, ^2–8,29^ it is not known whether mosquito salivary gland biology is under circadian control nor whether this influences the daily biology of sporozoites. To investigate this, we probed the transcriptome of salivary glands from infected female *Anopheles stephensi* mosquitos kept under 12h light/12h dark schedule (LD), or in constant dark (DD) for three consecutive days (Fig.1A). We used RNA-sequencing to assess the salivary gland transcriptome at different times of the 24h day and investigate the transcriptional changes associated with daily biological functions of the mosquito. We found that 27-49% of salivary gland genes displayed a cyclic expression profile, (Fig. 1B-E) representing 5-10x more than what has been previously reported for the mosquito’s head or body ^28^. Our discovery of a high proportion of cycling genes was further supported by hierarchical clustering analysis, which clustered all daytime time points (light or subjective day) based on genome-wide expression (Fig. 1C-D; S1A). Most circadian clocks are light entrained, and this is true for different mosquito species ^31,32^. A key feature of circadian clocks is that they persist in a cycle with approximately 24h period in the absence of environmental changes ^33,34^. Genes in the salivary glands of infected mosquitos cycled independently of the mosquito light/dark schedule (LD vs DD), suggesting that they are under circadian control and that the time of the day (rather than light) is the main driver for transcriptional fluctuations in mosquito salivary glands (Fig. 1E-G; S1A, C). By running permutation tests, we confirmed that the cycling genes identified are more than those that would be identified by chance (Fig. S2). Together, these results reveal substantial daily changes within mosquito salivary glands to potentially facilitate timing of a successful bloodmeal.

**Figure 1:**
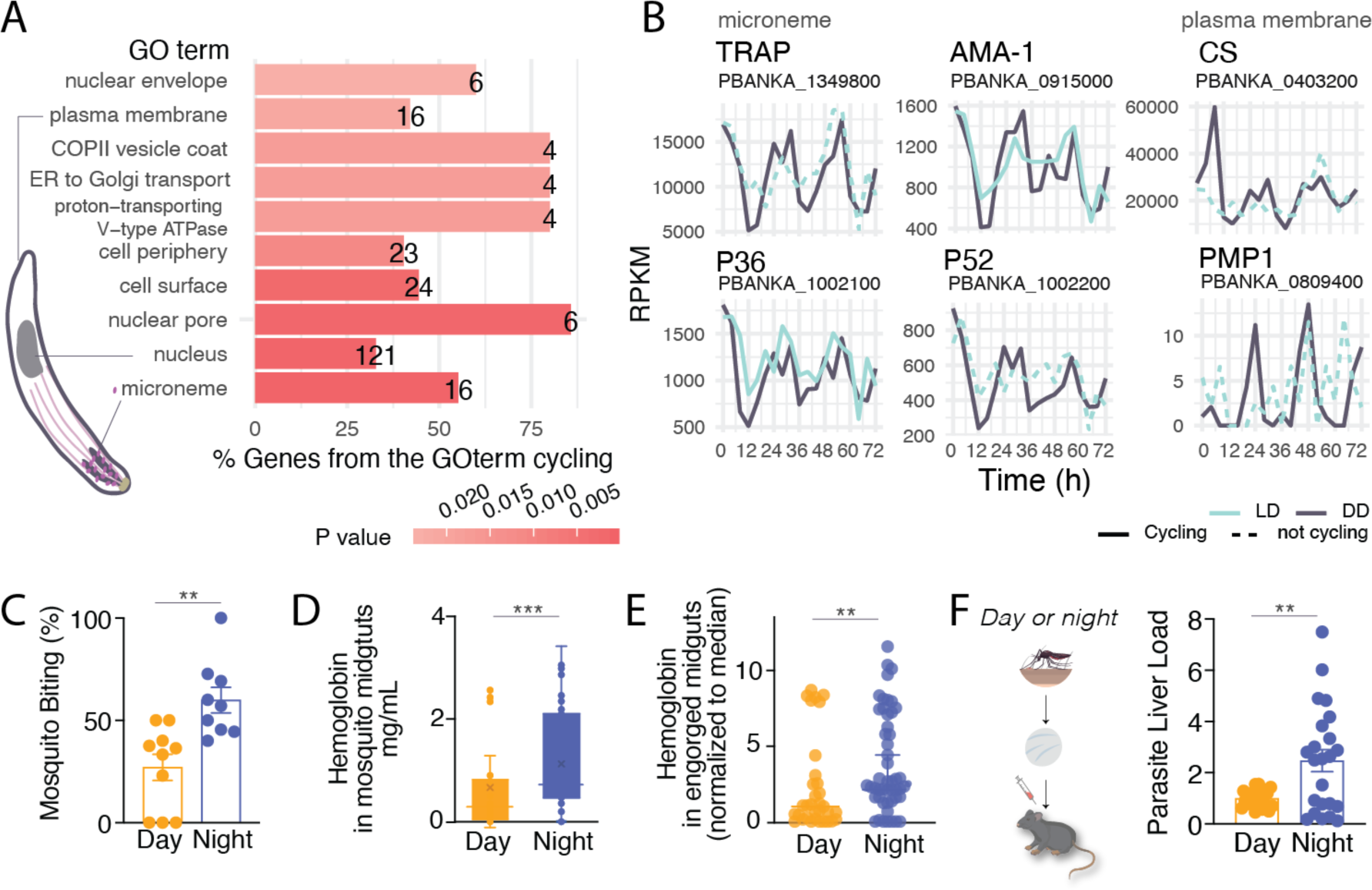
Half of the salivary gland transcriptome is rhythmic. **A.** Experimental design from two independent experiments for the sequencing of salivary glands from *Plasmodium berghei*-infected *Anopheles stephensi* mosquitos. Infected mosquitos were maintained in cyclic conditions (12h light / 12h dark). 18 days after mosquito infection, mosquitos were segregated into light/dark (LD) and constant dark (DD) conditions. 24h later, mosquitos were dissected every 4 hours over three days (72h) (n=32 samples for each light/dark (LD) and dark/dark (DD) condition). **B.** Representation of three genes from the top 20 cycling genes (based on significance) from each condition (Table S1-3). Orange and red lines represent the fit of two algorithms used to detect circadian oscillations (ARSER and JTK_cycle). Black lines represent the expression profile of salivary glands from infected mosquitos held in constant darkness (DD) and gray lines represent the 12h light / 12h dark condition (LD). **C-D.** Hierarchical clustering and heatmaps for each dataset by reordering of the timepoints according to similarity in gene expression. **E.** Heatmap of cycling genes from salivary glands from mosquitos in light/dark and dark/dark. Each row represents a gene. Gene expression is z-scored. **F.** Distribution of the peak of expression for all cycling genes in LD and DD conditions. This shows two main times of the day where most of the cyclic transcription occurs. Most gene expression peaks at 4h after light turn on, or beginning of daytime, and 4h after lights off, meaning 16h after beginning of daytime. **G.** The phase (time of peak of expression) for each of the 2,360 common genes is largely maintained across conditions (LD and DD). The plot represents a double plot of the phase of each cycling gene in each condition. **H.** Circadian fold-change in expression of the common cycling genes across conditions.

Overall, salivary gland genes reach peak expression early in the day and early in the night (Fig. 1F-G), about 4h later than what was previously reported for the whole mosquito head ^28^. This ‘dual rush hour’ of circadian gene expression is well described in other organisms, such as mammals and parasites ^25,35^. The 2,360 common genes that cycled in both LD and DD conditions had median oscillations of 3-fold amplitude and maintained their phases across conditions (Fig. 1H). Notably, some genes cycle with a circadian fold change above 100, such as arrestin-1 (ASTE007929) and guanine nucleotide-binding protein (ASTE009768).

The cycling genes in the salivary glands include the well-known circadian clock genes *Clock*, *Cycle*, *Period*, and *Vrille* (Fig. 1B and S3A), as well as genes that perform other biological functions (Fig. S3B-C), such as *Hyp15,* which encodes an *Anopheline*-specific protein that is highly enriched in adult female salivary glands ^36^ (Fig. 1B and Fig. S4). In fact, most of the mosquito transcripts that encode for bloodmeal-specific proteins cycled throughout the day (Fig. 2A-B and Fig. S3). These include the vasodilator *Peroxidase 5B* ^37^, the putative anti-inflammatory *D7 long form L2* (D7L2) ^38–40^, the anti-platelet aggregation *aegyptin*, anti-coagulant *anophelin/cEF*, and a gene of unknown function, *Salivary Gene 2* (SG2) precursor ^41^. When expression of these genes is knocked down, the efficiency of the mosquito bloodmeal has been shown to decline ^41^. Additionally, genes encoding SG1 family proteins, such as SAG (Saglin), TRIO (triple functional domain protein), and GILT (mosquito gamma-interferon-inducible lysosomal thiol reductase), that are associated with infection by malaria parasites ^18–20^ also showed a cycling profile (Fig. 2A). Saglin binds the parasite microneme TRAP (Thrombospondin Related Anonymous Protein) protein and is relevant for parasite invasion of the mosquito midgut and presence in the salivary glands ^20,42,43^. Immunization against the *Anopheles* TRIO protein can reduce the parasite burden in the host after mosquito transmission, suggesting this protein is also important for parasitic infection ^21^. GILT is a gamma interferon-inducible thiol reductase present in the saliva of infected mosquitos that reduces *Plasmodium* sporozoite cell traversal and transmission ^19^. Only female mosquitos take bloodmeals as they require specific nutrients (including proteins) from the blood to produce eggs ^44^, but both males and females can survive by feeding only on plant nectar. In addition to bloodmeal-associated salivary gland genes, we found that the glycolysis and gluconeogenesis pathways were also significantly enriched among cycling genes in LD and DD, with 9/11 genes of the glycolysis pathway cycling and with maximum expression in the first 4h of the day (Fig. 2C-F). Proteomics analysis comparing the levels of expressed proteins in the salivary glands in the morning (“Zeitgeber Time” ZT0) and evening (ZT12), showed 47 proteins differentially expressed across these time points (Fig. 2G-H and Fig. S5). These differentially expressed proteins include the two pyruvate dehydrogenase E1α and E1β of the glycolysis pathway (Fig. 2D and 2G-H, Table S6). Overall, these data suggest that mosquito salivary glands have gene expression and protein abundance rhythms that correlate with the mosquitos’ feeding behavior, in preparation for either a bloodmeal or sugar metabolism.

**Figure 2:**
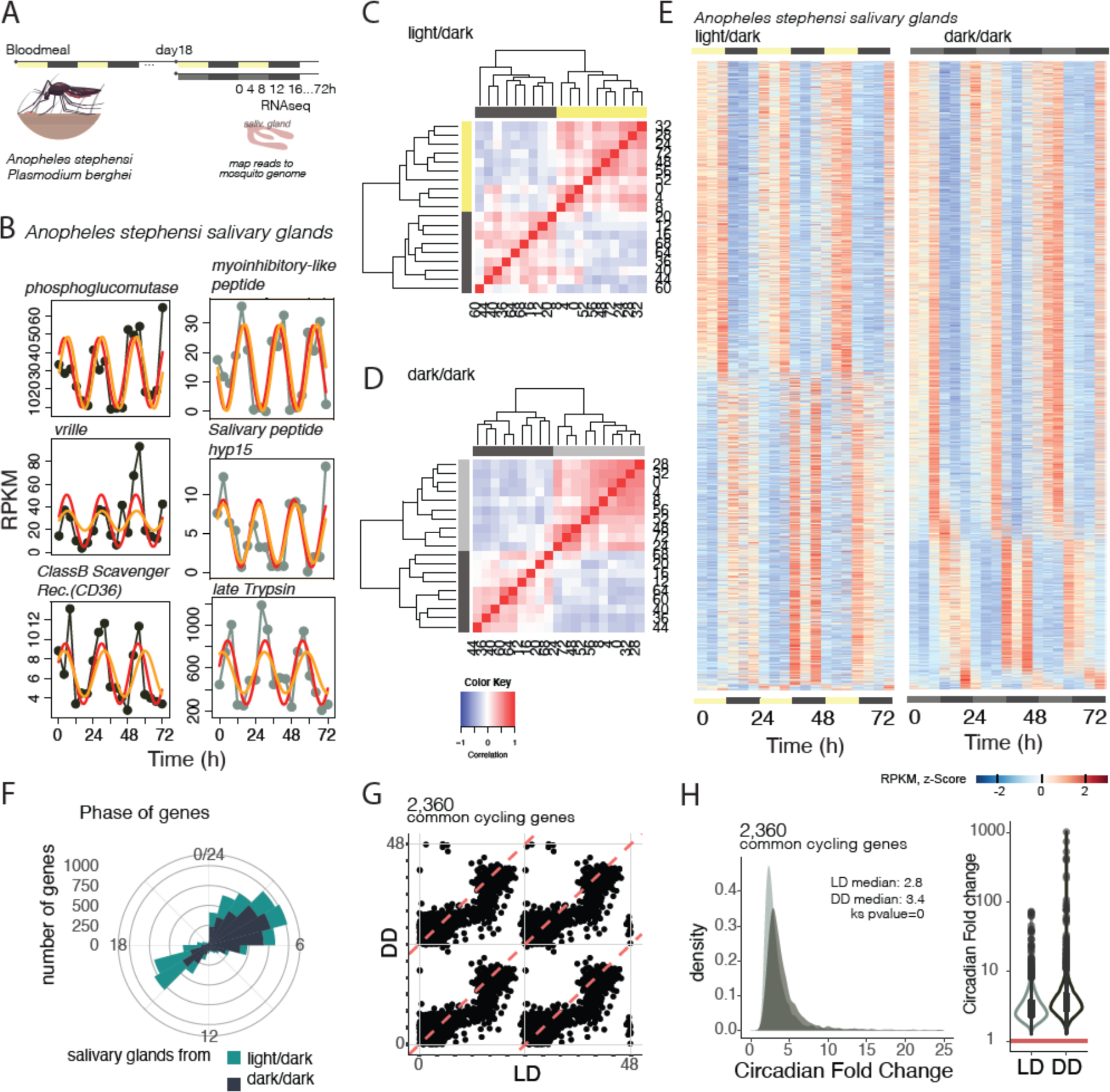
Salivary gland-specific transcripts cycle throughout the day. **A.** Rhythmic expression profile of nine well-characterized salivary gland proteins known to have a role in bloodmeal efficiency and/or malaria transmission (*Saglin, TRIO and GILT*). **B.** Expression profile of 21 transcripts encoding for proteins associated with blood feeding, identified by being differentially expressed between female and male salivary glands ^45^. 19 of these genes show similar phase and profile of gene expression. **C.** Top significant biological functions cycling under both LD and DD conditions at specific times of the day. Representative cycling genes from the pathway are from the DD condition. **D.** Representation of the Glycolysis and Gluconeogenesis pathway with the genes that code for the enzymes inside rectangular boxes. Only 2 out of the 11 represented genes of the glycolysis pathway do not cycle in the mosquito salivary glands throughout the day (unfilled boxes), whereas the other 9 cycle (colored boxes). **E-F.** Normalized expression profile (fold-change to mean RPKM expression values across time points) for each gene from the DD condition belonging to the Glycolysis and Gluconeogenesis pathway that cycle with a maximum expression between 0-4h, and from Pentose and Glucuronate interconversions between 4-8h. **G-H.** Protein expression levels from salivary glands across two time points, one in the morning (ZT0) and one in the evening (ZT12). Each point represents a pool of 5 salivary glands. Differential expression analysis in Fig. S5 and Table S6 with significance testing and fold-change calculation using MSstats. Mann Whitney test, * p < 0.05, ** p < 0.001, two-tailed).

### Sporozoites exhibit rhythmic gene expression within salivary glands

The transcriptome of sporozoites has been probed extensively ^46–50^. This parasite stage is often referred to as quiescent since sporozoites are thought to be cell cycle arrested and are transcriptionally similar across many days while in the salivary glands ^46,47,50^. Recent single cell RNA sequencing studies demonstrated heterogeneity within this population ^46,48^. Here, by probing the rodent parasite *P. berghei* sporozoite’s transcriptome at a very high temporal resolution during the daily cycle (Fig. 3A), we showed that 12-20% of sporozoite transcripts have cyclic expression (Fig. 3B and F). By running permutation tests, we confirmed that the cycling genes identified are more than those that would be identified by chance (Fig. S6). Additionally, sporozoite genes that cycle during the day have a maximum expression early in the morning (Fig. 3C-D), with an approximate median 3-fold circadian change in expression and a consistent phase (*i.e.*, peak expression time) in both light conditions (Fig. 3E).

**Fig. 3.**
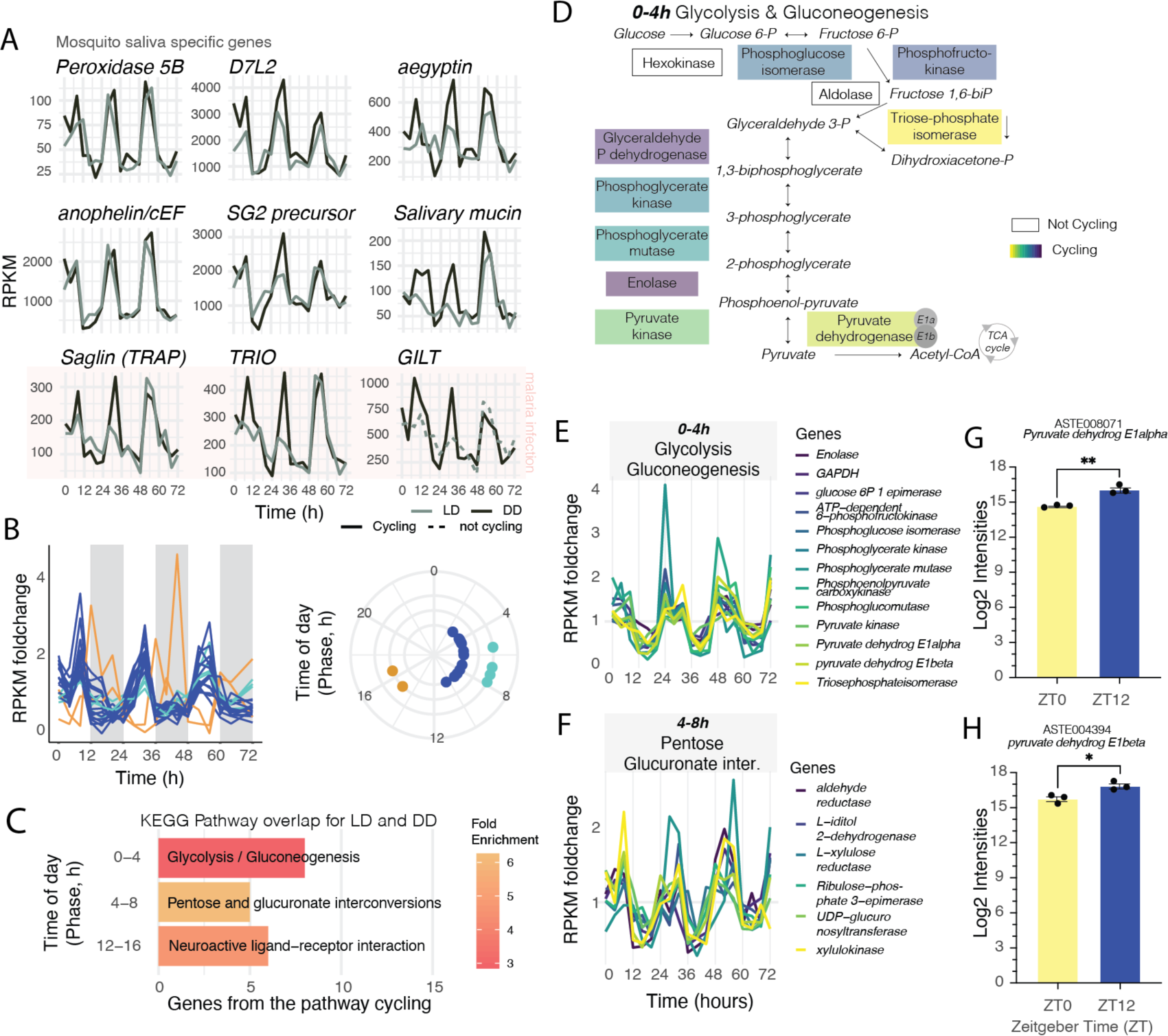
Transmission-associated genes from ‘quiescent’ sporozoites have daily rhythms. **A.** Experimental design from two independent experiments for the sequencing of *Plasmodium berghei* sporozoites from *Anopheles stephensi* mosquito salivary glands. Salivary glands were collected every 4 hours over three days for light/dark (LD) and dark/dark (DD) conditions. **B.** Representation of three cycling genes selected from the top 20 cycling genes (based on significance) for each condition (LD and DD) (see also Table S1, S4-5). Orange and red lines are the fitted curve of two of the algorithms used to detect circadian oscillations (ARSER and JTK_cycle). Gray and light green lines represent the profile of expression of each gene. **C.** Daytime distribution of the peak expression of cycling genes showing that cycling transcription peaks between 0-4h after lights on (beginning of daytime, light phase). **D.** Phase (time of peak of expression) for each of the 409 common genes (genes that cycle in LD and DD) is maintained in both conditions. **E.** Circadian fold-change of the common cycling genes in both conditions. **F**. Heatmap of sporozoite cycling genes from mosquitos in light/dark (LD) and dark/dark (DD) conditions. Each row represents a gene. Gene expression is z-scored. **G**. Representative immunofluorescence micrographs from purified salivary gland sporozoites at day 22-24 after mosquito’s bloodmeal at different timepoints (“Zeitgeber Time”) ZT3 (3h after lights on), ZT9, ZT15 and ZT21. DNA is shown in blue and circumsporozoite protein (CSP) is shown in red. Scale 5µm. **H**. Representative immunofluorescence micrographs of *Anopheles* mosquito midguts with oocysts from *P. berghei*-TK-GFP and GFP (parental) line after EdU incorporation. White circles delineate three oocysts. Arrowheads show oocysts that in the 48 hours prior to fixation, replicated their DNA as demonstrated by the incorporation of EdU. Asterisk* shows replicating mosquito midgut cells that incorporate EdU in the DNA. Scale 10µm. **I**. Representative immunofluorescence micrographs of salivary gland sporozoites from *P.berghei*-TK-GFP and GFP (parental) parasite line upon feeding of mosquitos with EdU. White line delineates a nucleus of a sporozoite. Sporozoites show no DNA replication as observed by the absence of EdU (red) at nuclear DNA (blue). Scale 5µm.

Once in the skin, sporozoite motility is crucial to cross the dermis and enter the blood or lymphatic systems ^51^. To reach and invade their unique target, the hepatocyte, sporozoites utilize substrate-dependent gliding motility and cell traversal ^52–56^. A multitude of cellular components, such as plasma-membrane proteins of the TRAP family, actin filaments, actin/myosin-based motor proteins, and the microneme, a specialized organelle from the Apicomplexa phylum ^56^, are critical for sporozoite motility ^57^. Strikingly, myosin A and thrombospondin-related sporozoite protein (TRSP) are among the top 20 cycling sporozoite genes (Fig. 3B and Table S1, S4-5). Other such genes include the transcription factor with AP2 domain and plasmepsin X, an aspartic protease important for invasion and egress during blood-stage infection, but whose function in sporozoites is unknown ^58,59^. Phospholipase gene expression is also rhythmic (Fig. 3B), and phospholipases have been shown to be involved in the migration of sporozoites through cells ^60^.

The consensus is that salivary gland sporozoites are non-dividing parasite forms. Nonetheless, due to technical challenges, the regulation of *Plasmodium* cell cycle has been understudied. Thus, we investigated whether the rhythmic gene expression we observed was the result of undocumented fluctuations in sporozoite numbers either due to i) migration of sporozoites into the salivary glands at specific times of day, or ii) unreported sporozoite replication within this tissue. To test whether cycling gene expression is independent of the numbers of sporozoites in the salivary glands, we normalized parasite reads by downsampling to the lowest number reads obtained in a timepoint and reanalyzed for daily oscillations. We found similar results when analyzing the total mapped reads or the downsampled reads (Fig. S7-8), suggesting that potential differences in parasite load across the day did not account for such daily gene expression changes.

Since *Plasmodium* parasites have rhythmic gene expression in the mammalian blood as they divide ^61–66^, we searched for evidence of cell division in salivary gland sporozoites throughout the day. We failed to identify sporozoites with morphologies suggestive of cell division, *e.g.*, duplicated nuclei and sporozoite separation (Fig. 3G) across different circadian timepoints. To eliminate the possibility that morphological alterations due to division would be a fast event that could be missed, we generated a transgenic *P. berghei* parasite line that expresses the thymidine kinase (TK) enzyme (*P. berghei*-GFP-TK). This parasite line allows the incorporation of the synthetic nucleoside analogue of thymidine, EdU (5-ethynyl-2′-deoxyuridine), into newly synthesized DNA, a method routinely used to detect and quantify DNA replication in mammalian cells. We provided EdU in the aqueous food of infected *Anopheles* mosquitos and analyzed its incorporation into parasites or mosquito nucleus. The EdU was available for 2 or 3 days to determine if cell division occurred during that time span. As expected, on day 13 post-bloodmeal, the DNA of the dividing stage of the parasite (oocysts) and dividing midgut cells of the mosquito, showed incorporation of EdU (Fig. 3H). In contrast, no EdU was incorporated in the nucleus of the sporozoites liberated from mosquito salivary glands 22 days post-bloodmeal, confirming that these are indeed non-replicating parasite forms (Fig. 3I).

Taken together, our results show that the transcriptional daily rhythms identified in sporozoites are not a consequence of cell division, but instead resemble a robust circadian rhythm. Despite being in a non-dividing stage, sporozoites are transcriptionally active parasite forms, with daily rhythms of gene expression that potentially allow parasites to anticipate timing of mosquito bloodmeal and prepare for an efficient transmission.

### Daily rhythms in sporozoite transmission

During a bloodmeal, few parasites are injected in the mammalian host, making this an ideal stage for the use of prophylactic drugs and vaccines ^67^. When we performed a functional analysis of all sporozoite cycling transcripts, the enriched GO terms whose genes cycle both under LD and DD conditions involved microneme and plasma membrane genes (Fig. 4A), with the myosin complex among the top 10 of GO terms in LD (Fig. S8C; S9B-C and Table S7). Over 50% of microneme-associated genes of the parasite cycle throughout the day. This includes the apical membrane antigen 1 (AMA1) and TRAP proteins (Fig. 4B), whose mutants have impaired motility ^53,57^ and whose antigenicity is being explored as vaccine candidates or monoclonal antibody treatment. Other microneme transcripts, whose protein function is essential to invade and transmigrate to the host cell also cycle ^68^. The circumsporozoite protein (CSP), the target of the first malaria vaccine (RTS,S) ^69,70^, is essential for the initial step of liver invasion by binding to highly sulfated proteoglycans in the liver sinusoids ^71,72^ and also shows a cyclic expression (Fig. 4B). Thus, we show that sporozoites have rhythmic transcription of motility-associated genes important for invasion and infection of the mammalian host.

**Fig. 4.**
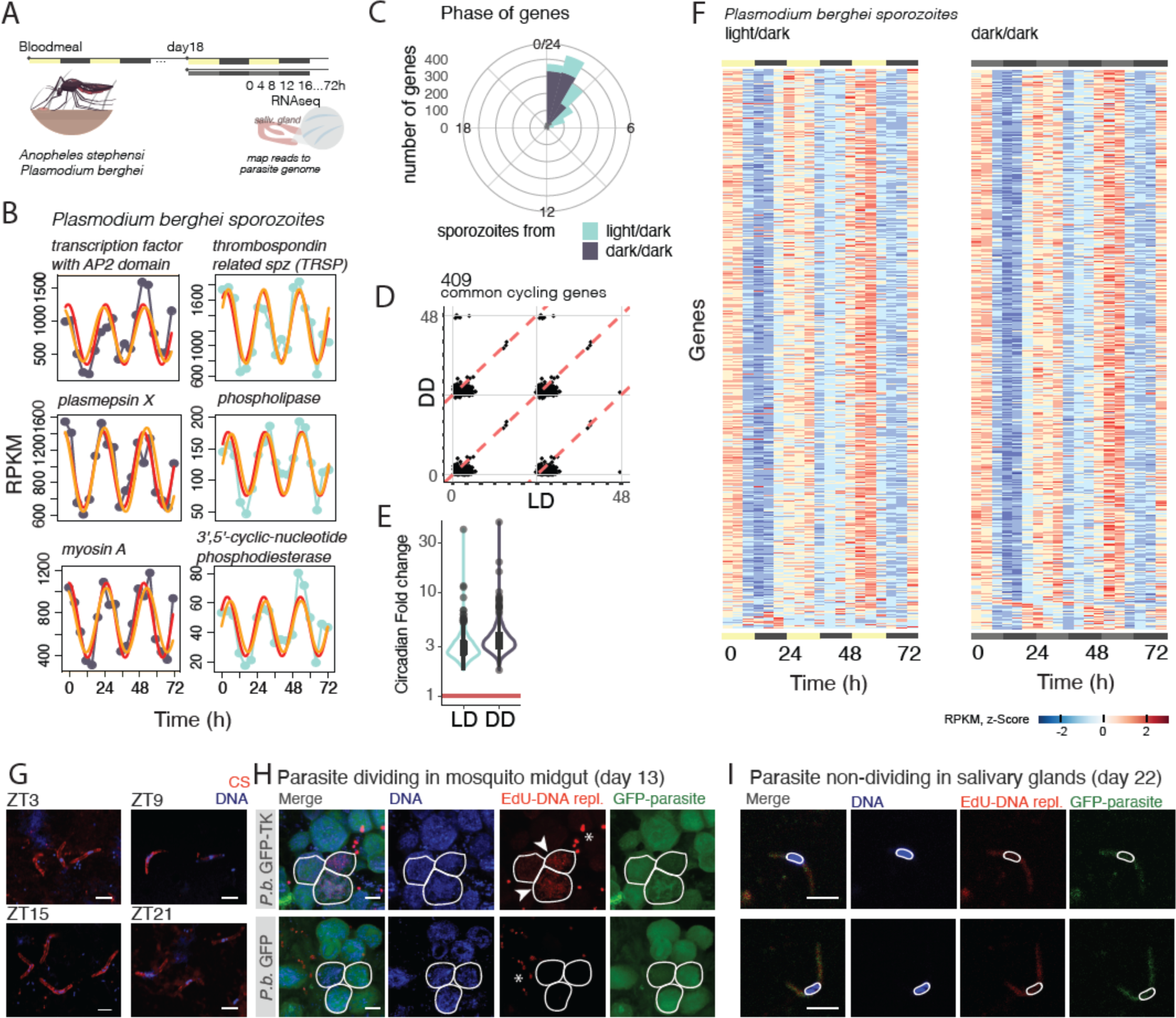
Daily rhythms are present in motility-associated genes and infection efficiency. **A.** The most significant biological functions with specific genes that have a cycling expression profile in DD. The same GO term functions cycle under LD conditions. Schematic of a sporozoite and some of its cellular compartments. **B**. Rhythmic expression profile of six microneme and plasma membrane-associated transcripts. **C.** Quantification of mosquitos that ensured a successful bloodmeal at ZT4 (daytime) and ZT16 (nighttime) as measured by the percentage of mosquitos that ingested blood within holding cups (n = 10 cups per group; n= 6-16 mosquitos within each cup; Mann Whitney test, ** p < 0.005. two-tailed). **D.** Quantification of hemoglobin in each mosquito midgut, as a proxy of blood ingested, after mosquito access to a bloodmeal for 30min (n = 67-68 midguts per group; 4 independent experiments; Mann Whitney test, *** p < 0.001, two-tailed). **E.** Quantification of hemoglobin in each engorged mosquito midgut, as a proxy of blood ingested, after mosquito access to a bloodmeal for 30min (n = 33-50 engorged midguts per group; 4 independent experiments; Mann Whitney test, ** p < 0.005, two-tailed). **F**. Quantification of parasite liver load in infected mice when the infection was initiated with intradermal injection of sporozoites dissected from mosquito salivary glands at ZT4 (daytime) and ZT16 (nighttime). Parasite load in the liver was assessed by qPCR normalized against *Hprt* (n = 21-23 mice per group; 5 independent experiments Mann Whitney test, ** p < 0.005, two-tailed).

In the blood-stage, malaria parasite has robust daily rhythms in gene expression ^61–65^; and these rhythms are host independent ^66^. However, there is an approximately 11h+ delay between the peak of mRNA transcription and protein expression ^73,74^. Although knowledge about the dynamics of transcription-translation for malaria parasites *in vivo* and for the sporozoite-stage is still lacking, we can hypothesize the existence of a complex regulatory mechanism for protein translation. In fact, highly transcribed mRNAs during the sporozoite stage, such as *UIS3* and *UIS4* ^75^, have undetectable protein content until later on in the parasite life cycle ^76,77^. However, when cross-referencing a dataset of sporozoite proteins expressed in the salivary gland parasites ^78^ with our cycling mRNA data, we found that 1/3 of the genes that cycle at the mRNA level are also translated into proteins in the salivary gland stage (Fig. S8-S10 and Table S7). These include myosin A, myosin B, myosin F, thrombospondin-related sporozoite protein, TRAP-like proteins, and AMA-1. These findings suggest that some of the sporozoite mRNAs that exhibit daily rhythms are translated into protein.

Since *Anopheles* mosquitos are more likely to bite at night and many of the salivary gland-specific genes are under circadian control (Fig. 1 and 2), this suggests that mosquitos are prepared to have a successful bloodmeal at specific times of the day. Indeed, we show an increase in successful bloodmeals when mosquitos bite at nighttime (Fig. 4C and Fig. S11A). Furthermore, we compared how much blood was ingested by quantifying hemoglobin levels inside the mosquito’s midgut after being allowed to feed. We observed a higher blood load in the midguts of mosquitos that fed at night (“Zeitgeber Time” ZT16, Fig. 4D and Fig. S11B, C). To test if the time of day also influenced the amount of blood ingested, we compared hemoglobin levels excluding the midguts that did not contain blood that at the time of dissection. We found that of those mosquitos that effectively ingested blood, the ones that fed during the nighttime had higher hemoglobin levels in their midguts (Fig. 4E and Fig. S11B-D), suggesting that, even if mosquitos can feed at any time, they are more likely to feed during the evening as their ability to successfully feed is also oscillating throughout the day. These behavioral and physiological changes across the day may have created an evolutionary pressure for sporozoites to be equipped with daily rhythms for a predictable encounter with a mammalian host at a specific time of the day. Possibly due to mosquito behavioral rhythms, when infected mosquitos are allowed to feed on mice during the morning or evening, there is usually higher parasite load in animals at nighttime (Fig. S11E). To test whether there are daily rhythms in transmission efficiency beyond mosquito biting, we dissected the salivary glands of infected mosquitos at the beginning of the day (ZT4) and the beginning of the night (ZT16) and infected mice with sporozoites by intradermal injection. We found a reduced parasite load in the livers of mice when infection was initiated during the daytime (daytime sporozoites and mice), compared to nighttime infection (nighttime sporozoites and mice) when mice showed a higher parasite liver load resulting from a more efficient transmission (Fig. 4D). This increase in the parasite load was abolished when sporozoites and mice time were mismatched (Fig. S11F). Taken together, our results suggest that the mammalian host, the *Plasmodium* parasite, and the *Anopheles* mosquito vector evolved to coordinate daily rhythms, ensuring a successful transmission that occurs at a specific time of the day.

## Discussion

In conclusion, we have uncovered a previously unrecognized level of mosquito-parasite interaction, revealing that sporozoites are transcriptionally active parasite forms with cyclic RNA expression, equipped to cope with host-seeking behavior and ensure efficient infection. Our results lead us to propose that sporozoites have evolved to be rhythmic to cope with (or exploit) the rhythms of vector and/or host. The daily transcriptional rhythms of sporozoites suggest that they are ‘poised’ or ‘primed’ for their crucial task of encountering and infecting the mammalian liver at a specific time, which aligns with the mosquito’s feeding time. Altogether, our study suggests that the rhythms of mosquito, parasite *and* host must be aligned for a successful malaria transmission. Further studies that dissect the contributions of each of the steps and clock mechanisms for the observed phenomenon will be needed. Currently, our findings highlight the importance of timing in mosquito biting and establishing new host infections, contributing to our understanding of malaria transmission biology.

We have observed almost half of the mosquito transcriptome cycling within their salivary glands. Such a high proportion of cycling genes, when compared to previous studies ^28^, could be explained by the increased sensitivity and statistical power of circadian algorithms, resulting from a longer time course at high sample frequency and thus resolution in our study. Additionally, since previous analyses were performed from whole mosquito body parts, potential conflicting rhythms of different tissues could flatten the overall oscillations, which would lead to an underestimation in the number of 24h-cycling transcripts. Alternatively, the expression of salivary gland genes may be more rhythmic than that observed in the head and body. Here we focused on infected *Anopheles* mosquitos; however, since many of the cycling genes encode for proteins that regulate fundamental cellular functions of cells such as glycolysis, it is possible that these observations are extendable to uninfected mosquitos, female or male. In addition, further characterization of mosquito circadian rhythms, including these of the salivary glands should be done across the lifespan of the mosquito as rhythms may decline as mosquito age, similar to what was demonstrated in mammals ^79^, and across multiple bloodmeals.

In the future, it will be critical to identify the molecules responsible for signaling the time-of-day information to the mosquito salivary glands and sporozoites and explore whether these are intrinsic circadian rhythms. This knowledge could help explore methods to disrupt the synchronization between mosquito vectors and malaria parasites, possibly through disrupting the biological clocks of either the mosquitos or the parasites to prevent efficient transmission. Moreover, further research is warranted by the potential impact of the use of bed nets on changing mosquito biting behavior regarding new infections and parasite biology ^80^. Strategies that leverage the knowledge of peak mosquito activity and bloodfeeding times could be considered to optimize the timing of insecticide application or other vector control measures. For example, targeting insecticide spraying during periods of increased mosquito activity could maximize effectiveness. These findings of daily rhythms in disease transmission are likely widespread to other vector borne diseases.

Overall, our high temporal resolution study provides valuable insights into the intricate interactions between mosquitos, malaria parasites and mammalian hosts, contributing to the development of novel intervention strategies against malaria transmission. By recognizing the active nature of sporozoites and their preparation for infection, we advance our understanding of malaria biology and open new avenues for combatting this devastating infectious disease.

## Materials and Methods

### Parasite lines

Parasites used in this study: GFP expressing *P. berghei* ANKA (clone 259cl2, *Pb*-GFP); *P. berghei* ANKA resistance marker free GFP expressing line (clone 440cl1, obtained from the Leiden Malaria Research Group, http://www.pberghei.eu, ^81^ and *P. berghei* ANKA-GFP line expressing the thymidine kinase (TK) from *Herpes simplex* virus (*Pb*-GFP-TK).

The *Pb*-GFP-TK parasite line was generated by double-homologous recombination targeting the P230p locus of the GFP expressing *P. berghei* ANKA clone 440cl1. The recombination construct containing the P230p locus sequence flanking the expression cassette for the TK and the human DHFR coding sequences fused via a 2A self-cleaving peptide was kindly provided by Kathrin Witmer (Imperial College, London). Transfection was performed in blood stage merozoites (as described by^81^). Briefly, blood from a *Pb*-clone 440cl1 infected BALB/c mouse was collected and cultured for 16 hours *in vitro*. Mature schizonts were purified by a Nycodenz gradient and transfected with the recombination construct using the Amaxa electroporation system (Lonza). Transfected merozoites were injected into the tail vein of a BALB/c mouse (6–8 weeks of age) and selected by the administration of Pyrimethamine in the mice drinking water (70μg/ml). Pyrimethamine resistant parasite population containing the correct genomic integration of TK/DHFR expressing cassette was cloned by limiting dilution and injection of one parasite per mouse (10 BALB/c male mice, 6–8 weeks of age). Genomic DNA was isolated from the blood of infected animals (NZY Blood DNA isolation kit) and the successful recombination at the modified locus was verified by PCR (NZYTech PCR mix) with the following oligonucleotide pairs: for the TK coding sequence CAC TTG ACA GAA TGG CCT CGT; CTG CAG ATA CCG CAC CGT AT) and genomic integration (GCA AAG TGA AGT TCA AAT ATG; GGG CAT TTT CTG CTC CGG GC).

### Mosquito rearing and infections

*Anopheles stephensi* mosquitos were reared at 28°C and 80% humidity under a 12-h/ 12-h light/dark cycle and fed on 10% glucose (Sigma) and 0.2% 4-aminobenzoic acid (Millipore Sigma) in MilliQ water changed daily. Female *Anopheles stephensi* mosquitos 4-5 days old were allowed to bite two mice infected with *Plasmodium berghei* ANKA (GFP_con_, 259cl2 for one hour). For the study of parasite DNA replication, mosquitos were infected with *Plasmodium berghei* ANKA-GFP-TK. After the infectious bloodmeal, mosquitos were kept in 12h light/12h dark, at constant temperature of 20°C. Mosquitos were fed by cotton pads soaked with 10% glucose (Sigma) and 0.2% 4-aminobenzoic acid (Millipore Sigma) in MilliQ water changed daily.

### Mice

Mice were maintained under specific pathogen-free conditions and housed at 23°C with a 12h light/12h dark schedule in accordance with the European regulations concerning Federation for Laboratory Animal Science Associations, category B and the standards of the University of California Berkeley Institutional Care and Use Committee. C57BL/6J were purchased from Charles River or Jackson Laboratories.

### DNA replication analysis

Parasite’s DNA replication was assessed by analysing EdU incorporation in *Plasmodium berghei* ANKA-GFP-TK (thymidine kinase). The parental line, *Plasmodium berghei* ANKA-GFP (clone 440cl1) was also subjected to the same protocol.

Infected mosquitos were fed for 48h or 72h with 200 μM of EdU (Invitrogen) in mosquito’s food. Salivary glands from 20 infected mosquitos were then collected into basal DMEM (Gibco) medium on days 22-24 post-bloodmeal every 6h for 48h (timepoints: ZT0 (at lights on), ZT6, ZT12 (at lights off) and ZT18). Salivary glands were grounded with a plastic pestle and filtered through a 40 mm Falcon cell-strainer (Thermo Fisher Scientific) to release sporozoites. Sporozoites were purified using accudenz column purification method ^82^. Then, 20 μl of purified sporozoites were allowed to adhere to the wells of a glass slide (Thermo Scientific ER-308B-CE24, 10 wells 6.7 mm glass) and fixed for 10 minutes with 4% PFA. Sporozoites were then washed 2x in 3% BSA for 10 minutes, permeabilized for 20 minutes in 0.5% Triton-X100 in PBS and then washed twice for 10 minutes in 3% BSA. Click-iT chemistry reaction mix was prepared according to the manufacturer’s instructions (Invitrogen, Click-It EdU Imaging Kits, C10339) by addition of a fluorescent azide through a Cu(I)-catalyzed reaction. Sporozoites were washed with 3% BSA in PBS, followed by a PBS wash for 10 minutes, incubated overnight at 4°C with anti–UIS4 (goat, 1:500, from Sicgene) and washed 3 times with PBS. The following secondary antibodies: anti-GFP rabbit polyclonal antibody conjugated to Alexa Fluor 488 and donkey anti-goat Alexa Fluor 568 (all 1:1000 from Life Technologies, Invitrogen) were incubated for 1 hour at RT in 3% BSA. Cell nuclei were stained with Hoechst 33342 in 1:1000 (from Life Technologies, Invitrogen Invitrogen). Sporozoites were then washed 3× in PBS and slides mounted using Fluoromount-G (SouthernBiotech). All images were acquired on Zeiss confocal microscope (LSM 880) with 63× amplification and processed in ImageJ.

Two independent positive controls for actively dividing parasites (oocysts) were obtained by dissecting midguts from 20 mosquitos fed with EdU at days 13-14 after a blood meal, followed by fixation with 4% PFA for 20 minutes at room temperature (RT). Midguts were then washed 2x in 3% BSA for 10 minutes and permeabilized for 1 hour in 0.5% Triton-X100 in 3% BSA and then washed twice for 10 minutes in 3% BSA. Click chemistry reaction mix was prepared according to the manufacturer’s instructions (Invitrogen) and added to the midguts for 30 minutes at RT. These were then washed with 3% BSA and incubated overnight at 4°C with anti–*P. berghei* HSP70 (2E6, 1:300) followed by 3 washes with 3% BSA and incubation for 2 hours with secondary antibodies at RT. The following fluorescently tagged secondary antibodies were used for detection: anti-GFP rabbit polyclonal antibody conjugated to Alexa Fluor 488 and goat anti-mouse conjugated to Alexa Fluor 647 (all 1:1000 from Life Technologies, Invitrogen). In parallel, cell nuclei were stained with Hoechst 33342 in 1:1000 (from Life Technologies, Invitrogen Invitrogen). Then midguts were washed 3× in PBS and placed a coverslip on top of the slide mounted using Fluoromount-G (SouthernBiotech). All images were acquired from 20 dissected midguts using the Zeiss confocal microscope (LSM 880) with 63× amplification and processed in ImageJ.

### Salivary gland tissue collection for RNA sequencing

Two independent experiments were done. On day 16 post-bloodmeal, mosquitos were divided into smaller cages holding ∼30-40 mosquitos. On day 19, mosquitos were either maintained in the 12h light/dark schedule (LD) or kept in constant darkness (DD) for salivary gland collection that started on day 20 every 4h for 2 days (experiment 1) or on day 21 for 3 days (experiment 2). When combining both experiments, for each dataset (LD or DD) the first 48h (13 timepoints) have two independent experiments, each timepoint with a pool of salivary glands from ∼20 mosquitos. The additional 24h (6 timepoints) have one replicate of pooled salivary glands (∼20 mosquitos). Each dissected mosquito showed ∼30-50,000 sporozoites/mosquito. Mosquito euthanasia was achieved by CO2 to avoid a putative cold shock response of gene expression. Dissections of salivary glands were performed in PBS with RNAse inhibitor (Invitrogen) (0.1U/μl) to avoid a nutrient rich environment that could reset the circadian rhythm of cells and each dissected tissue was snap frozen with a minimum volume of PBS. RNA was extracted from whole salivary glands using TRIzol LS according to manufacturer’s instructions (Life Technologies). Measurements were taken from the samples discussed in this paragraph.

### Transcriptome sequencing (RNAseq) and data analysis

Total RNA (1 - 2.5 μg) was enriched for mRNA using 50 μl of Poly(A) beads for RNAseq according to the manufacturer’s instructions (Invitrogen). The integrity of RNA and removal of ribosomal RNAs was confirmed on a TapeStation (Agilent Technologies). Sequencing libraries were constructed using the TruSeq RNA Sample preparation protocol (Illumina). Quantification of library concentration was achieved by Qubit fluorometric quantitation (Thermo Fisher Scientific) and size of library calculated by TapeStation. RNA-sequencing of libraries was performed on the Hiseq2000 (Illumina) with 50 bp reads for the first experiment 0-48h LD and DD, and on the NextSeq 550 (Illumina) with 75 bp reads for the second experiment 0-72h LD and DD, according to the manufacturer’s instructions by the UTSW McDermott Next Generation Sequencing Core. Read quality was assessed using the FASTQC quality control tool (http://www.bioinformatics.babraham.ac.uk/projects/fastqc). Reads were mapped with STAR ^83^ following Cutadapt trimming ^84^ to the following genomes: PlasmoDB-26_PbergheiANKA from www.PlasmoDB.org and Anopheles-stephensi-SDA-500_AsteS1.6 from www.vectorbase.org. In the infected salivary glands, an average of 11% of the uniquely identified reads mapped to the parasite. Due to being evolutionarily distant, only 0-1% of the uniquely mapped reads mapped to both genomes. The number of reads mapping to each gene was determined and then normalized to RPKM (reads per kilobase of transcript per million mapped reads).

In this study we generated two independent datasets for infected salivary glands in LD and DD conditions. The results presented are an average of both experiments but analyzing the individual experiments provided similar observations (Fig. S1). To exclude that the variability between pooled samples/time points could influence circadian analysis due to different numbers of sporozoite reads sequenced in each sample*, i.e.*, how infected each pool of salivary glands were, we ran the downstream analysis with both total uniquely mapped reads and with the same datasets downsampled to the lowest uniquely mapped reads. This approach allowed us to eliminate fluctuation of read number mapping to either species across timepoints. Both analyses showed the same results (Fig. S7 and S8) further supporting that the daily rhythms observed in gene expression of both salivary glands and sporozoites was not a consequence of infection load variation in each timepoint.

For further quality control of salivary gland dissection, the obtained gene expression from these datasets (LD and DD) was compared to the defined fat body transcriptome. With VectorBase.org, fat body genes were considered fat-body-specific if genes had a fold change of 10 or greater compared to other tissues (adult female fat body sample and the reference samples were adult female Malpighian tubules, midgut, and ovary) ^85^. The result was 489 protein-coding genes with a FC >= 10. Out of these 489 up-regulated fat body genes, 443 genes were shown to be expressed in our dataset. To determine relative expression of these 443 up-regulated fat body genes in our dataset, the mean gene expressions of all genes across all timepoints in the LD and DD salivary gland sample (our dataset) were normalized by the max gene expression value of the same dataset. Similarly, all expressed genes with a TPM value > 1 in the VectorBase dataset were normalized to their max gene expression. This analysis further supported that the dataset we analyzed is salivary gland specific (Fig. S2B). Downstream bioinformatic analysis after RPKM calculations was performed in RStudio. Hierarchical clustering and heatmaps were obtained for each dataset by reordering the timepoints according to gene expression using function heatmap.2 from the *gplots* package ^86^ (Fig. S1 and S7). Both hierarchical clustering analysis and heatmaps of Spearman correlations were performed on centered log2-transformed RPKM values.

### Time-series analysis for circadian cycling

A gene was considered expressed if its mean expression, across all timepoints and conditions, was higher than 0.1 RPKM for the mosquito and higher than 0.5 RPKM for the parasite. With this definition, 11,383 genes, 83.6% of the mosquito genome, were identified as expressed in salivary glands; from sporozoites in salivary glands, 4,670 of the genes, 88.9% of the parasite genome, were expressed. Cycling of mRNA was assessed using four circadian statistical algorithms within the MetaCycle package ^87^, which implements Lomb-scargle, JTK_CYCLE ^88^, ARSER ^89^, and the RAIN methods ^90^. A gene was considered cycling if two of the four programs detected significant periodic expression with a threshold of P ≤ 0.05, and that in one of them an FDR adjusted P value ≤ 0.05. Phase reported by ARSER was used for further analyses. Full table of all expressed genes and their cycling analysis is reported for each species and condition in Tables S2-S5. Circadian fold-change was calculated by dividing the mean maximum expression across the three days of collection by the mean minimum expression across those days.

### Permutation test

To determine the number of genes that were identified as cycling by chance, we analyzed 1000 permutations of all cycling genes given by the re-ordering of the sampled timepoints of collection and its corresponding *Plasmodium* and *Anopheles* gene expression in LD as well as DD. Using the MetaCycle R package, each permutation was analyzed for cycling genes using the JTK, LS, and ARSER algorithms. Each gene was labelled as cycling or non-cycling based on the combined q-value of all 3 algorithms using Fisher’s method and defined a cycling gene with a combined q-value of <= 0.05, two-tailed.

### Functional analysis

To identify the genes that cycle at a transcriptional level, gene orthologs from *Anopheles gambiae* were obtained using Biomart from www.vectorbase.org. Overall, the functions of mosquito salivary cycling genes were analyzed by KEGG pathway enrichment and InterPro protein domain enrichment using DAVID ^91^. The top 10 most significant molecular pathways sorted by fold enrichment in both environmental conditions (LD and DD) are represented (Fig. S4) and reported in detail in Table S2. To acquire a time-of-day resolution of functions, mosquito cycling genes were divided into 4h resolution phase groups and analyzed for KEGG pathway and InterPro using DAVID. The most significant pathway or protein domain enriched in both environmental conditions (LD and DD) are reported for each of the 4h intervals. No significant enrichment overlap among conditions was found for 8-12h, 16-20h or 20-24h for KEGG pathways (Fig. 2). *Plasmodium berghei* functional analysis was assessed for all cycling genes due to apparent peak expression in the first 4h of the day, in both conditions (LD and DD). GO term for biological process and KEGG pathway analysis were performed in www.PlasmoDB.org (Fig. 4 and S9) and reported in detail in Table S7. By overlapping our dataset of cycling mRNAs with the Lindner 2013 dataset of sporozoite proteins expressed in the salivary gland ^78^, we identified 238 genes that cycle at the mRNA level and whose proteins are expressed during sporozoite stage in the salivary glands (Fig. S8D and Table S6).

### Proteomics analysis

**Protein extraction**. Infected salivary glands were collected from female *A. stephensi* mosquitos and snap frozen. For each timepoint (ZT0 and ZT12), 3 replicates were collected where each replicate had 5 pairs of infected salivary glands. 50 μL of fresh RIPA protease inhibitor cocktail was added to each sample pellet, Piece RIPA Buffer (Thermoscientific, Cat No: 23225 and 23227), 1x Halt^TM^ Protease & Phosphatase Inhibitor Cocktail (100x) (Thermoscientific, Ref No: 1861281) and 1x 0.5 M EDTA Solution (100x) (Thermoscientific, Ref No: 1861274) and samples homogenized with micropestles. Samples were then vortexed and incubated on ice for 10 minutes to facilitate lysis. The salivary gland homogenate suspension sample was spun in a microcentrifuge for 20 minutes for 12,000 rpm at 4 °C. The protein supernatant was transferred, and protein quantified using the Pierce**^TM^** BCA Protein Assay Kit (Thermoscientific, Cat No: 23225 and 23227). Preparation of diluted standards and sample preparations were done according to manufacturer’s instructions. Briefly, each protein lysate sample was diluted at 1:2 in the RIPA protease inhibitor cocktail. 200 μL working reagent was added to each sample well and shaken for 30 minutes with a plate shaker. The 96-well plate was then incubated for 30 minutes at RT. Absorbance was measured at 562 nm using the CLARIOstar plate reader. A linear regression equation and standard curve were calculated from the absorbance values of these diluted standard duplicates using PRISM.

**Peptide preparation with S-Trap micro columns.** A solution of Tris (pH 8) and SDS in water was added to sample lysates to achieve a final concentration of 5% sodium dodecyl sulfate (SDS), 50mM Tris in 57.5 μL total sample volume. For reduction and alkylation of cysteines, the samples were incubated at 37°C, 1200 rpm with 5mM dithiothreitol (DTT; Pierce) for 1 hour, then 14 mM iodoacetamide (IAA; Pierce) for 45 min. Subsequent binding and washing of the samples on S-Trap micro columns (Protifi) were performed following the manufacturer’s protocol. For protein digestion, 50mM tetraethylammonium bromide (TEAB) containing trypsin (Pierce) and Lys-C (FUJIFILM Wako Pure Chemical Corporation) was added to the S-trap column and incubated at 47°C for 1.5 hrs. Each protease was used at a protease to protein ratio of 1:20 (w/w) or minimum of 0.5μg. Digested peptides were sequentially eluted with 50mM TEAB, 0.2% formic acid (FA), and 50% acetonitrile (ACN). The resulting eluates were dried in a SpeedVac and resuspended in 0.1% FA, 0.015% n-dodecyl-β-D-maltoside (DDM). Peptide concentrations were measured by NanoDrop, and the samples were further diluted in 0.1% FA, 0.015% DDM to a final concentration of 10ng/μL.

**LC-MS/MS data acquisition.** Samples were analyzed on a timsTOF SCP (Bruker Daltonics) coupled to an EvoSep One liquid chromatography system (Evosep Biosystems). Samples were prepared for injection by loading 20μg of peptides each onto Evotip Pures (Evosep Biosystems). Peptides were eluted online from the Evotips and separated by reverse-flow chromatography on a 15 cm PepSep column (75 μm internal diameter, 1.9 μm C18 beads; Bruker Daltonics) with a 10 μm zero-dead volume sprayer (Bruker Daltonics), using the Whisper 20 method (60-min gradient length, 100 nl/min flow rate) from Evosep. The column was maintained at 50°C in a Bruker column toaster.

Data was acquired in dda-PASEF mode (data-dependent acquisition with parallel accumulation serial fragmentation) with high sensitivity detection enabled. Ions were delivered to the timsTOF SCP through CaptiveSpray ionization and analyzed across a mass range of 100-1700 m/z and mobility range of 0.7-1.3 1/k0. Accumulation and ramp time were both set to 166 ms, with a 100% duty cycle. For each MS1 scan, 5 MS2 PASEF ramps were performed for a total cycle time of 1.03 ms. A polygon filter was used to exclude singly charged precursor species. Quadrupole isolation of precursors for MS2 fragmentation was set to allow a 2 m/z window for precursors under 700 m/z and a 3 m/z window for precursors above 800 m/z. Collision energy linearly increased with ion mobility, from 20eV at 0.6 1/k0 to 65eV at 1.6 1/k0. The intensity threshold for precursor repetitions was set to 500, with a target intensity of 20,000. Active exclusion of precursors was released after 0.2 min.

**Raw data searching and data analysis.** The raw LC-MS/MS data was searched with Fragpipe 20.0 against a combined *Anopheles stephensi* SDA-500 (www.vectorbase.org) and *Plasmodium berghei* ANKA database (www.PlasmoDB.org), with contaminant and reverse sequences added by Fragpipe. Redundant protein sequences were consolidated into single FASTA entries. Entries were reformatted to allow Fragpipe to parse out relevant information, including Protein ID and description, organism, and gene name. Default parameters for closed, tryptic search were used, with the inclusion of deamidation (NQ) and phosphorylation (STY) as additional variable modifications. MS1 quantification was also performed, without match-between-runs and normalization across runs enabled. The MSstats.csv output generated by Fragpipe was used for downstream analyses. Features mapping to contaminant proteins were removed prior to data processing with the MSstats R package. The dataProcess function was run with default parameters, which performs log2 intensity transformation, normalization by equalizing medians, run-level protein intensity summarization, and imputation using an accelerated failure time model. Significance testing and fold-change calculation between conditions was performed using the MSstats function groupComparison (Table S6).

### Quantification of mosquitos’ blood meal

**Mosquito biting**. Mosquitos were reared in two incubators kept in opposite light/dark cycles. Mosquitos were transferred to small cups containing between 5 and 20 mosquitos each and fasted for 12h prior to the biting experiment. Mosquitos were allowed to feed on either an anesthetized mouse or through membrane feeding on human blood for these experiments. When allowed to bite on anesthetized infected mice at two times of the day – at ZT4 (4h after lights on) and at Z16 (4h after lights off) – mosquitos had access to the mouse for 30 minutes in their respective light settings. For membrane feeding, the glytube protocol was adapted ^92^. For assembling the heating element of the glytube, heat-resistance plastic sealing film was soldered onto the 50mL conical tube to create a seal and prevent glycerol leakage. 2 layers of stretch parafilm 5 cm x 5 cm were stretched over the heat-resistance plastic sealing film. Glytubes were prewarmed overnight or for a minimum of 30 minutes before the feeding experiment at 50°C. 1mL washed human blood (red blood cells, BioIVT) was prewarmed for 15-30 mins at 37°C and placed onto the cap reservoir with stretched parafilm for mosquito feeding. The inverted glytubes were held in place on top of the cage mesh by a metal clamp stand for mosquitos to feed for 30 minutes in their respective light settings.

Successful mosquitos bloodmeal was accessed by observing the mosquito abdomen for enlargement and presence of blood during midgut dissection using a light microscope. To obtain % mosquito biting, the total number of mosquitos with a bloodmeal was divided by the total number of mosquitos in the cup (an independent experiment). The behavioral assay was performed 6 independent times for both daytime and nighttime timepoints.

**Mosquito blood ingestion**. Midguts of each mosquito were dissected into 100 μL HBSS (Gibco) and immediately frozen. On analysis day, samples were thawed and homogenized with a pestle. Samples were spun at 5,400 rpm for 8 minutes and the supernatant collected to a 96-well plate and hemoglobin colorimetry assessment performed as recommended by the manufacturer (Invitrogen). In brief, diluted high sensitivity standards were prepared and the samples were incubated with the diluent for 30 mins at room temperature. Hemoglobin colorimetry was measured using plate reader absorbance at 562 nm at room temperature. A standard curve using 2 replicates of raw diluted standard absorbance values was calculated using a linear regression equation generated from Prism and/or by the MARS software from the ClarioStar plate reader. The linear regression equation was used to calculate the hemoglobin concentration (mg/mL) from raw sample absorbance values. ZT4 and ZT16 feeding experiments done on the same day were analyzed together in the hemoglobin colorimetry assay, where each time a new standard curve was generated from diluted standards via linear regression analysis. Sample hemoglobin concentrations were calculated from a linear regression equation generated from each standard curve.

### Parasite load in mouse liver

*Anopheles stephensi* mosquitos infected with *Plasmodium berghei* ANKA-GFP (259cl2) were maintained in 12h light/12h dark cyclic conditions.

From mosquito bite. Mice were anesthetized with 90mg/kg of ketamine and 5mg/kg of xylazine. Each mouse was placed ventral side down on a mesh-covered enclosure containing 15-20 female *Anopheles stephensi* mosquitos. At ZT4 (4h after lights on, when mosquitos do not typically bite) and at Z16 (4h after lights off, close to natural biting time of *A. stephensi*) 21 days after mosquito’s blood meal, mosquitos were allowed to feed freely for 25 minutes at the appropriate light, temperature and humidity to the mosquito time of day. Livers were collected 46 hours after the midpoint of infection and snap frozen. The whole tissue was homogenized in 4 mL of TRIzol (Thermo Fisher Scientific), ¼ processed according to manufacturer instructions. cDNA synthesis was performed with 500ng of purified RNA using the UltraScript cDNA Synthesis Kit (Genessee ScientificNZYTech), as per manufacturer’s instructions. cDNA was then used for quantitative Polymerase Chain Reaction (qPCR) using PowerTrack SYBR Green (ThermoFisher) with the pair of primers for mouse glyceraldehyde-3-phosphate dehydrogenase (*mGapdh*: CAA GGA GTA AGA AAC CCT GGA CC; CGA GTT GGG ATA GGG CCT CT) and parasite 18S ribosomal RNA (*Pb18S rRNA*: AAG CAT TAA ATA AAG CGA ATA CAT CCT TAC; GGA GAT TGG TTT TGA CGT TTA TGT G). Measurements of SYBR fluorescence were performed on BioRad CFX96 C1000 Real-Time PCR Systems (Bio-Rad Laboratories) and analyzed using the 2^-ΔΔCt^ method.

From sporozoite injection. 21 days after mosquito’s blood meal, to ensure the sporozoites were fully mature, salivary glands from ∼50 infected mosquitos at ZT4 (4h after lights on, when mosquitos do not typically bite and close to the peak of sporozoite gene expression) and at Z16 (4h after lights off, close to natural biting time of *A. stephensi*) were collected into basal DMEM (Gibco) medium. Salivary glands were ground with a plastic pestle and filtered through a 40 mm Falcon cell-strainer (Thermo Fisher Scientific) to release sporozoites. Sporozoites were then counted using a hemocytometer (Marienfeld Superior, Lauda-Königshofen, Germany) followed by intradermal injection in the left hind leg of each mouse with 30,000 sporozoites in 50 µl of DMEM. We designed two experimental groups: Light - injection of ZT4 sporozoites in 5 mice from light environment; and Dark – injection of ZT16 sporozoites in 5 mice from dark environment, recreating what normally happens in the field. Two additional groups were mismatched, meaning that sporozoites collected from ZT4 mosquitos were injected in mice at the opposite ZT, *i.e.*, ZT16, and the complementary group. 32 hours after injection, livers were collected and homogenized in 3ml of TRIzol (Thermo Fisher Scientific) containing 0.1 mm Zirconia/Silica beads (BiospecTM Products, Bartlesville, OK, USA). Liver homogenization was performed through mechanical disruption in a Mini-BeadBeater (BioSpec Products) for 1.5 min. The 3mL of homogenized tissue were divided into 3 clean 1.5 mL microcentrifuge tubes (Eppendorf) each with 1 mL of tissue lysate. One aliquot of 1 mL was used for extraction of total RNA by addition of 200µl of chloroform, followed by a 4°C centrifugation at 12,000g for 15 minutes. The upper aqueous phase was then transferred to a new microcentrifuge tube and RNA extraction proceeded according to the manufacturer’s instructions of the NZY Total RNA Isolation Kit (NZYTech, Lisboa, Portugal). Quantification of total RNA concentration was performed on NanoDrop 2000 spectrophotometer (Thermo Fisher Scientific) according to the manufacturer’s guidelines. cDNA synthesis was performed with 1 µg of purified RNA using the NZY First-Strand cDNA Synthesis Kit (NZYTech), as per manufacturer’s instructions. cDNA was then used for quantitative Polymerase Chain Reaction (qPCR) using iTaq Universal SYBR Green Supermix (Bio-Rad Laboratories, Hercules, CA, USA) with the pair of primers for mouse Hypoxanthine-guanine phosphoribosyltransferase (*mHprt*: CAT TAT GCC GAG GAT TTG GA; AAT CCA GCA GGT CAG CAA AG) and parasite 18S ribosomal RNA (*Pb18S rRNA*: AAG CAT TAA ATA AAG CGA ATA CAT CCT TAC; GGA GAT TGG TTT TGA CGT TTA TGT G). Measurements of SYBR fluorescence were performed on ViiA 7 (384-well plates) Real-Time PCR Systems (Thermo Fisher Scientific).

## Acknowledgments

We thank Ana Parreira for maintaining the mosquito colony and establishing their infections, Izabela Kornblum for assistance preparing the sequencing libraries and Gokul Kilaru for assistance with downsampling. We thank the SporoCore at University of Georgia, Athens for providing mosquitos. We thank Vanessa Zuzarte-Luis, Ksenija Slavic, Maria Inês Marreiros, João Vieira and Viriato M’Bana for help dissecting mosquito salivary glands and Ângelo Chora and V. Zuzarte-Luis for the intradermal injections. We would like to thank Catherine Merrick, Camila Mariano and Kathrin Witmer for the help generating the *P. berghei*-GFP-TK parasite line.

## Data availability

All sequencing data are deposited in GEO datasets (www.ncbi.nlm.nih.gov/geo/query/xxxx). Accession codes will be available before publication.

